# Biophysical network models of phase-synchronization in MEG resting-state

**DOI:** 10.1101/2021.08.04.455014

**Authors:** N Williams, B Toselli, F Siebenhühner, S Palva, G Arnulfo, S Kaski, JM Palva

## Abstract

Magnetoencephalography (MEG) is used extensively to study functional connectivity (FC) networks of phase-synchronization, but the relationship of these networks to their biophysical substrates is poorly understood. Biophysical Network Models (BNMs) have been used to produce networks corresponding to MEG-derived networks of phase-synchronization, but the roles of inter-regional conduction delays, the structural connectome and dynamics of model of individual brain regions, in obtaining this correspondence remain unknown. In this study, we investigated the roles of conduction delays, the structural connectome, and dynamics of models of individual regions, in obtaining a correspondence between model-generated and MEG-derived networks between left-hemispheric regions. To do this, we compared three BNMs, respectively comprising Wilson-Cowan oscillators interacting with diffusion Magnetic Resonance Imaging (MRI)-based patterns of structural connections through zero delays, constant delays and distance-dependent delays respectively. For the BNM whose networks corresponded most closely to the MEG-derived network, we used comparisons against null models to identify specific features of each model component, *e*.*g*. the pattern of connections in the structure connectome, that contributed to the observed correspondence. The Wilson-Cowan zero delays model produced networks with a closer correspondence to the MEG-derived network than those produced by the constant delays model, and the same as those produced by the distance-dependent delays model. Hence, there is no evidence that including conduction delays improves the correspondence between model-generated and MEG-derived networks. Given this, we chose the Wilson-Cowan zero delays model for further investigation. Comparing the Wilson-Cowan zero delays model against null models revealed that both the pattern of structural connections and Wilson-Cowan oscillatory dynamics contribute to the correspondence between model-generated and MEG-derived networks. Our investigations yield insight into the roles of conduction delays, the structural connectome and dynamics of models of individual brain regions, in obtaining a correspondence between model-generated and MEG-derived networks. These findings result in a parsimonious BNM that produces networks corresponding closely to MEG-derived networks of phase-synchronization.

**Highlights:**

- Simple biophysical model produces close match (ρ=0.49) between model and MEG networks
- No evidence for conduction delays improving match between model and MEG networks
- Pattern of structural connections contributes to match between model and MEG networks
- Wilson-Cowan oscillatory dynamics contribute to match between model and MEG networks

## 1. Introduction

Functional Connectivity (FC) networks underpin the execution of cognitive tasks (Cole et al. (2013), Cohen & D’Esposito (2016)) and are also observed during resting-state (Beckmann et al. (2005), Damoiseaux et al. (2006)). FC networks are sets of interacting brain regions, wherein the interactions are reflected by correlated activities of the brain regions. Key forms of correlation observed in Electroencephalography (EEG) and Magnetoencephalography (MEG) resting-state or task data, are the pairwise correlations between the oscillation amplitudes of a set of brain regions (Hipp et al. (2012), Colclough et al. (2015), Brookes et al. (2012)), or the pairwise synchronization between oscillation phases of a set of brain regions (Palva et al. (2005), Palva et al. (2010), Palva & Palva (2012), Zhigalov et al. (2015), Siebenhühner et al. (2020), Marzetti et al. (2019)). FC networks of amplitude correlation or phase-synchronization are widely studied with EEG/MEG, but the structure-function mapping between biophysical substrates and FC networks is poorly understood. We refer to biophysical substrates as systems-level components, *e*.*g*., the structural connectome, dynamics of brain regions or inter-regional conduction delays, which underpin observed FC networks.

Biophysical Network Models (BNMs) (Woolrich & Stephan (2013)) have been used to relate MEG-derived networks of amplitude correlation to their biophysical substrates. BNMs comprise models of individual brain regions, linked by biologically informed structural connectivity. BNMs of Kuramoto oscillators linked by diffusion Magnetic Resonance Imaging (dMRI)-based structural connections with realistic conduction delays, produced FC networks of amplitude correlation corresponding moderately to those observed in MEG resting-state (Cabral et al. (2014)) (ρ = 0.41, where ρ is Pearson Correlation). Similarly, BNMs of spiking neuron populations linked by dMRI-based structural connections with realistic delays also produced FC networks of amplitude correlation corresponding moderately to those observed in MEG-derived networks of amplitude correlation (ρ = 0.4) (Nakagawa et al. (2014)).

FC networks of phase-synchronization have been suggested to be relevant to information processing in the brain, coordinating communication across regions via regulation of spike - time relationships (Fries (2005), Singer (1999), Palva & Palva (2012)) and supporting functional integration (Palva & Palva, (2012), Siegel et al. (2012), Deco et al. (2015)). In fact, particular phase-synchronization networks are known to be recruited for specific cognitive tasks, including working memory (Kitzbichler et al. (2011), Palva et al. (2010)), visual attention (Lobier et al. (2018)) and sensorimotor processing tasks (Hirvonen et al. (2018)).

Just as for MEG-derived networks of amplitude correlation, BNMs have been employed to relate EEG- or MEG-derived networks of phase-synchronization to their biophysical substrates. BNMs of Kuramoto oscillators linked by dMRI-based structural connections with conduction delays, produced networks of phase-synchronization corresponding to those observed in EEG resting-state (Finger et al. (2016)). However, this model produced networks with only weak correspondence to EEG-derived networks, when the networks were estimated with measures insensitive to EEG volume conduction (ρ = 0.17 for weighted Phase Lag Index (wPLI)). BNMs of Wilson-Cowan oscillators with inhibitory synaptic plasticity (ISP), linked by dMRI-based structural connections with conduction delays, have been used to produce networks of phase-synchronization corresponding to those observed in MEG resting-state (Abeysuriya et al. (2018)). While this BNM produced networks with a moderate correlation to MEG-derived networks (ρ = 0.43 for Phase Locking Value (PLV), ρ = 0.28 for Phase Lag Index (PLI)) (Abeysuriya et al. (2018)), the strengths of model-generated phase-synchronization (∼0 - 0.6 for PLV and ∼ 0 - 1 for PLI) were an order of magnitude higher than those observed in MEG-derived networks (∼0.01 - 0.06 for both PLV and PLI). Further, while the correspondence between the model-generated and MEG-derived networks was assessed, the roles of conduction delays, the structural connectome and the dynamics of models of individual brain regions in producing this correspondence was not investigated.

In this study, we investigated the roles of conduction delays, the structural connectome and the dynamics of models of individual regions, in producing networks corresponding to observed MEG-derived networks of phase-synchronization between left-hemispheric regions. To do this, we first compared three hypotheses, respectively postulating model-generated networks of phase-synchronization corresponding to MEG-derived networks, are produced by the dynamics of Wilson-Cowan oscillators linked by dMRI-based patterns of structural connections, via zero, constant and distance-dependent conduction delays. We expressed each of these hypotheses as a BNM. For the BNM producing networks corresponding most closely to the observed MEG-derived networks, we used comparisons against null models to determine specific features of each model component, for *e*.*g*., pattern of connections in the structural connectome, that contribute to the correspondence between the model-generated and MEG-derived networks. Together, these investigations yield insight into the precise roles of inter-regional conduction delays, the structural connectome and the dynamics of models of individual brain regions, in producing networks corresponding to observed MEG-derived networks of phase-synchronization.

## 2. Materials & Methods

In this study, we investigated the roles of conduction delays, the structural connectome and the dynamics of models of individual brain regions, in producing model-generated networks corresponding to the observed MEG-derived networks of phase-synchronization. To do this, we first compared the correspondence between model-generated and MEG-derived networks of three BNMs, respectively including zero, constant and distance-dependent conduction delays. Figure 1 illustrates the pipeline to estimate and compare the model-generated and MEG-derived networks. For the BNM producing networks corresponding most closely to the MEG-derived networks, we used null models to determine specific features of each model component that contributes to the correspondence between model-generated and MEG-derived networks. For example, we compared the correspondence between model-generated networks from the original model and the MEG-derived networks, against the correspondence between model -generated networks from null models comprising degree-preserved randomised versions of the structural connectome, and the MEG-derived networks. Such a comparison enabled inferences on the contribution of the *pattern* of structural connections to the correspondence between model-generated and MEG-derived networks, over and above the number of connections, *i*.*e*. degree, to each brain region.

**Figure 1.**
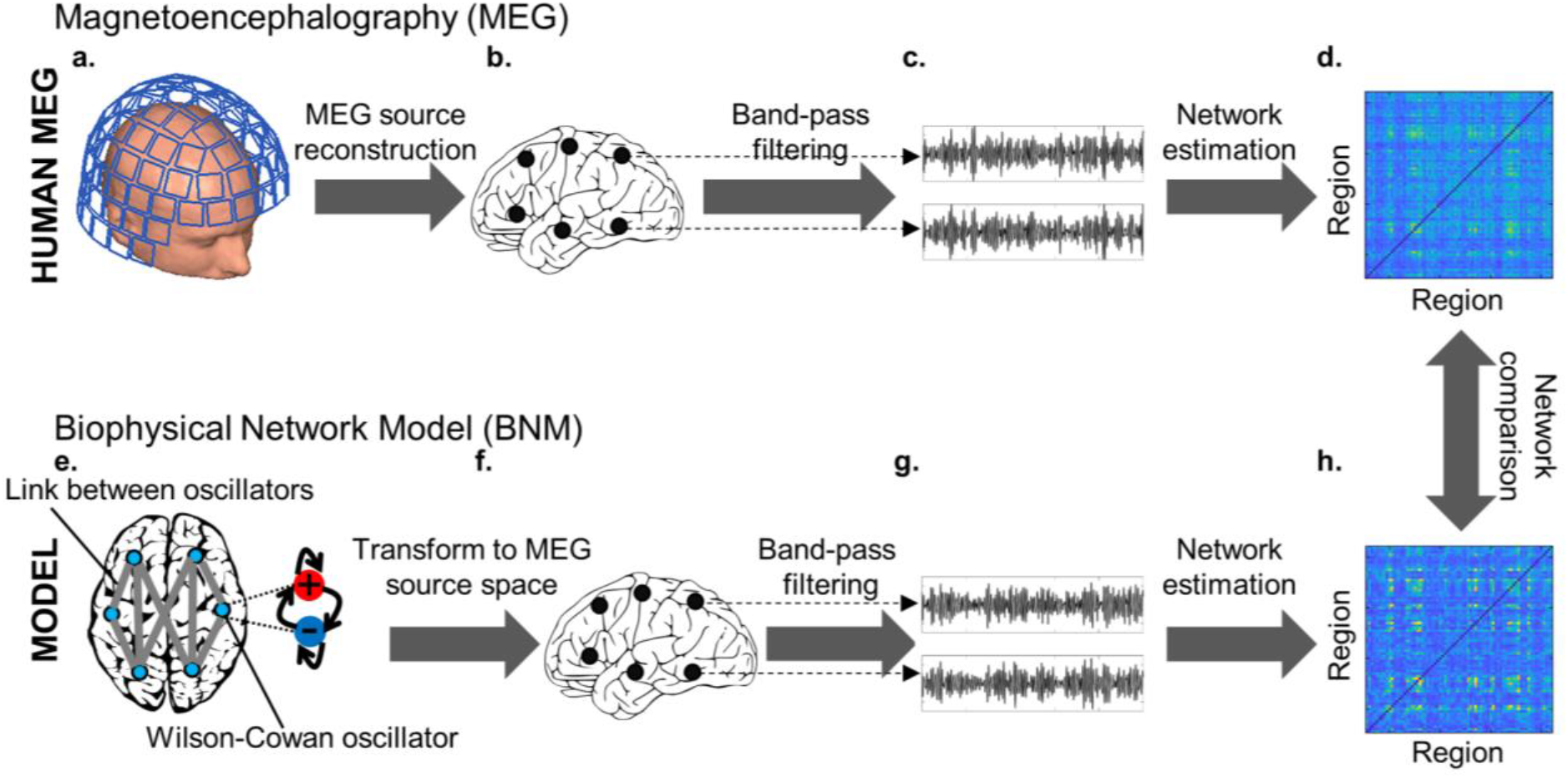
Pipeline to generate and compare model-generated and MEG-derived networks of phase-synchronization. a. MEG device with sensors detecting weak magnetic fields arising from post-synaptic potentials in the human brain. b. Source-reconstructed data projected from MEG sensors to brain regions. c. Band-pass filtered source-reconstructed MEG data. d. Matrix of strength of phase-synchronization between source-reconstructed MEG data from every pair of brain regions. e. Biophysical Network Model (BNM) comprising Wilson-Cowan oscillators linked by biologically informed structural connections. f. Simulated data transformed to MEG source space by projecting dynamics to MEG sensors, and projecting sensor-level simulated data back to brain regions. g. Band-pass filtered source-reconstructed simulated MEG data. h. Matrix of strength of phase-synchronization between source-reconstructed simulated MEG data from every pair of brain regions. MEG device panel from Pfeiffer et al. (2018).

### 2.1. Models

We implemented three models, each representing specific hypotheses on the interaction between dynamics of models of brain regions, structural connectome and conduction delays, in producing networks corresponding to MEG-derived networks of phase-synchronization.

#### Wilson-Cowan zero delays model

Wilson-Cowan oscillatory dynamics interact with the pattern of connections in the structural connectome without conduction delays, to produce networks of phase-synchronization corresponding to those observed in MEG resting-state.

We implemented this model as a set of Wilson-Cowan oscillators linked by biologically informed structural connectivity (Wilson & Cowan (1972), Kilpatrick (2013), Cowan (2016)). The dynamics of each oscillator arise from the interaction between excitatory and inhibitory neuronal populations, *i*.*e*. the PING model (Traub et al. (1997)), and are also influenced by external inputs and the dynamics of linked oscillators. For oscillator *i*:

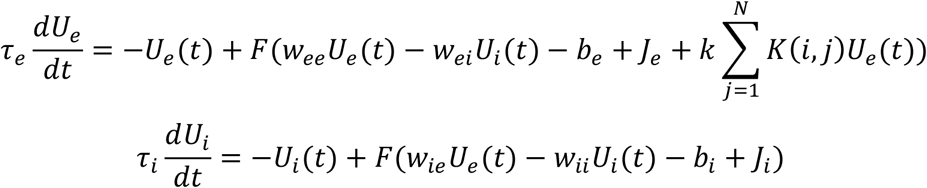

where 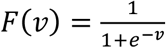 is a sigmoid function, *U*_*e*_ (*t*) and *U*_*i*_ (*t*) are the mean firing rates at time *t* of the excitatory and inhibitory populations respectively, *w*_*ee*_ and *w*_*ii*_ are the excitatory-excitatory and inhibitory-inhibitory connection weights respectively, *w*_*ie*_ and *w*_*ei*_ are the excitatory-inhibitory and inhibitory-excitatory connection weights, *b*_*e*_ and *b*_*i*_ are threshold constants for excitatory and inhibitory populations, *J*_*e*_ and *J*_*i*_ are injection currents to excitatory and inhibitory populations, *τ*_*e*_ and *τ*_*i*_ are the time constants of the excitatory and inhibitory populations, and *k* is a scalar multiplier over the coupling matrix *K*, which specifies links between *N* oscillators. For this model, we postulated that non-zero conduction delays are not relevant to producing the networks of phase-synchronization observed in MEG resting-state.

#### Parameter values

We set the parameters of each Wilson-Cowan oscillator to the same values (Table 1) since we assumed the intrinsic properties of the brain regions are identical. The choice of values for connection weights within (*w*_*ee*_, *w*_*ii*_) and between (*w*_*ie*_, *w*_*ei*_) excitatory and inhibitory populations, and the strengths of external currents (*J*_*e*_, *J*_*i*_) are consistent with canonical neurophysiological findings of 1.) strong excitatory-excitatory connections (Jansen & Rit (1995), Douglas et al. (1989), Binzegger et al. (2004)) 2.) excitatory-inhibitory and inhibitory-excitatory connections being weaker than excitatory-excitatory connections (Jansen & Rit (1995), Binzegger et al. (2004)) 3.) weak inhibitory-inhibitory connections (Douglas et al. (1989), Binzegger et al. (2004)) and 4.) the strength of injection currents, *e*.*g*. from thalamus, being significantly lower than strength of excitatory-excitatory connections (Douglas et al. (1989), Binzegger et al. (2004)). The threshold constants of excitatory (*b*_*e*_) and inhibitory populations (*b*_*i*_) are difficult to determine from the neurophysiology literature, but the values were chosen to produce oscillatory phenomena (Wilson & Cowan (1972), Singh et al. (2016), Sreenivasan et al. (2017)). We chose the excitatory (*τ*_*e*_) and inhibitory time constants (*τ*_*i*_) to produce dynamics around 10 Hz, *i*.*e*. the peak frequency of oscillatory power in the power spectrum of MEG resting-state (Nakagawa et al. (2014)). We tuned the parameter *k, i*.*e*. the scalar multiplier over the coupling matrix, *K* (see Section 2.6).

**Table 1.**
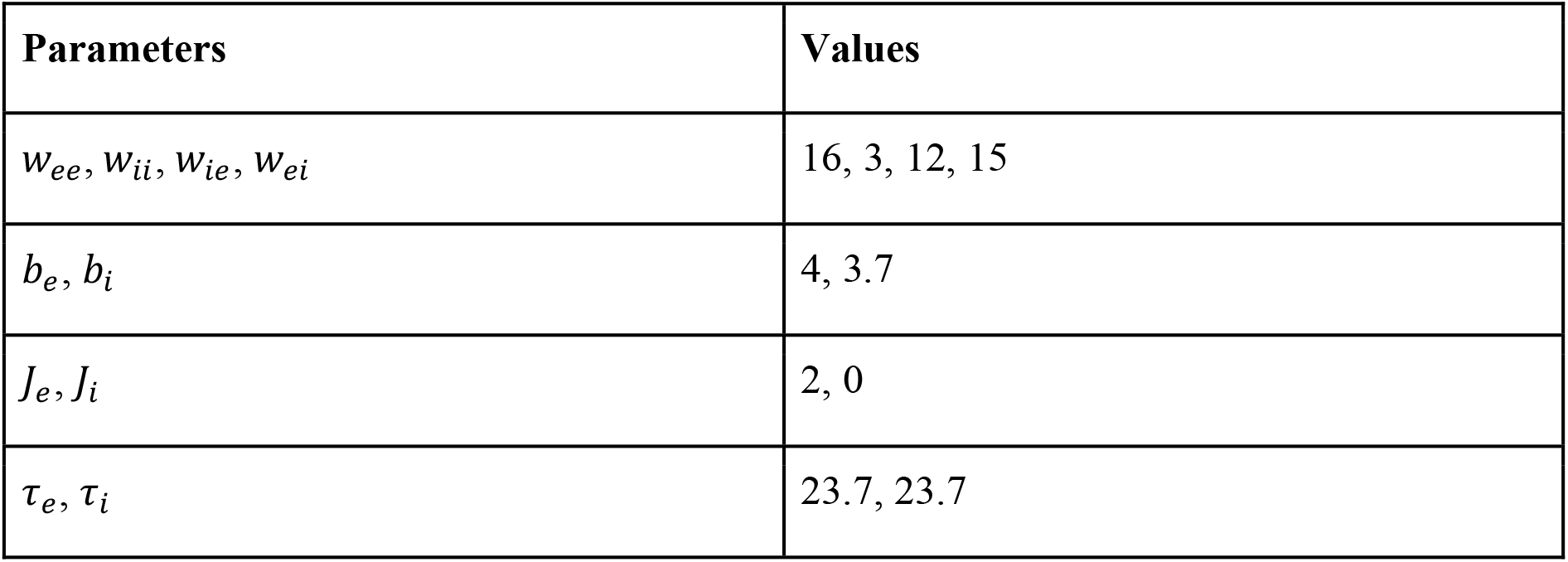
Parameter values of individual Wilson-Cowan oscillators

#### Wilson-Cowan constant delays model

Wilson-Cowan oscillatory dynamics interact with the pattern of connections in the structural connectome through constant non-zero conduction delays, to produce networks of phase-synchronization corresponding to those observed in MEG resting-state.

We implemented this model as a set of Wilson-Cowan oscillators linked by biologically informed structural connectivity with constant, non-zero delays. The dynamics of individual oscillators were influenced by inter-regional interactions of constant delay, in contrast to zero delays assumed by the Wilson-Cowan zero delays model. For oscillator *i*:

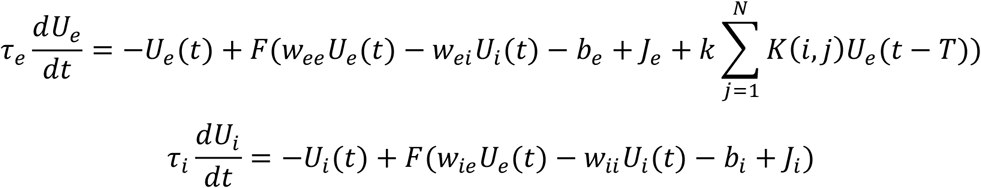

where *T* specifies the constant delay in milliseconds, and all other terms bear the same definitions as the Wilson-Cowan zero delays model. For this model, we postulated that the increase in conduction delay due to longer distance between regions is counteracted by the decrease in conduction delay due to higher conduction velocity between regions. This assumption is supported by recent modelling work (Noori et al. (2020)) reporting highly similar conduction delays across region-pairs due to increased conduction velocity produced by activity-dependent myelination counteracting longer delays between distant brain regions.

#### Parameters values

We set the parameters of the Wilson-Cowan oscillators to the values in Table 1, and tuned the parameters *k* and constant delay *T* (see Section 2.6).

#### Wilson-Cowan distance-dependent delays model

Wilson-Cowan oscillatory dynamics interact with the pattern of connections in the structural connectome through distance-dependent conduction delays, to produce networks of phase-synchronization corresponding to those observed in MEG resting-state.

We implemented the Wilson-Cowan distance-dependent delays model as comprising a set of Wilson-Cowan oscillators linked by biologically informed structural connectivity, with conduction delays proportional to distance between respective brain regions. The dynamics of individual oscillators are influenced by inter-regional interactions of variable delay, in contrast to zero and constant delays assumed by the Wilson-Cowan zero delays and constant delays models respectively. For oscillator *i*:

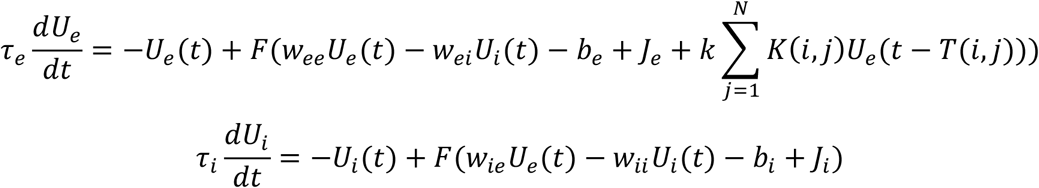

where *T*(*i, j*) specifies the distance-dependent delay in metres/second, and all other terms bear the same definitions as the model describing the Wilson-Cowan zero delays model. For this model, we specified conduction delays as Euclidean distance between regions divided by a conduction velocity *ν* (Cabral et al. (2014), Nakagawa et al. (2014), Hadida et al. (2018)).

Rather than Euclidean distance, an alternative method to estimate distances between brain regions is the average length of the streamlines between regions, as calculated by the tractography method used. However, we found the average estimated streamline lengths of ∼20% of structural connections to be lower than the corresponding Euclidean distances, which represent the minimum length between regions. Since this might be due to known biases in streamline termination with tractography methods (Smith et al. (2013)), we chose Euclidean distance to estimate distances between brain regions.

#### Parameters values

We set the parameters of the Wilson-Cowan oscillators to the values in Table 1, and tuned the parameters *k* and the conduction velocity *ν* (see Section 2.6).

### 2.2. Model comparison

We compared the Wilson-Cowan zero delays, constant delays and distance-dependent delays models by the correspondence between their respective networks of phase-synchronization and the observed MEG-derived network of phase-synchronization. We ran 20 simulations of each model to obtain its dynamics at different starting conditions. After each run, we compared the model-generated network of phase-synchronization to a held-out MEG-derived network of phase-synchronization, *i*.*e*. an MEG-derived network not used to estimate model parameters (see Section 2.5). We quantified correspondence between model-generated and MEG-derived networks, *i*.*e*. model performance, by the RMSE (Root Mean Square Error) and Pearson Correlation between the upper-triangular elements of the model-generated and MEG-derived adjacency matrices, low RMSE and high Correlation reflecting a close correspondence. We used independent samples *t*-tests of the respective RMSE and Correlation values to compare model performances, with two-tailed *p* < 0.001 considered statistically significant. Further, we used Cohen’s *d* (Cohen (1988)) to measure effect sizes, with *d* ≥ 1 considered a strong effect.

### 2.3 Obtaining model-generated network of phase-synchronization

Below, we describe the details of the different model components, model simulations, and the procedure to generate the network of phase-synchronization from the model dynamics.

#### 2.3.1 Model elements

The model comprised an ensemble of 74 Wilson-Cowan oscillators linked by biologically informed structural connectivity. We used the left-hemispheric regions of the Destrieux brain atlas (Destrieux et al. (2010)) to specify the number and location of the oscillators. We specified the links between oscillators by the binary matrix of the strongest 10 percentile structural connections between left-hemispheric regions of the Destrieux atlas. We derived this binary matrix from a group-averaged (N=57) weighted matrix of the number of streamlines between brain regions, estimated by constrained spherical deconvolution (Smith et al. (2013)) and probabilistic tractography (Smith et al. (2012)) on pre-processed DWI images from the Human Connectome Project (van Essen et al. (2013)). We divided each element of the binary matrix by the sum of elements in its row, to ensure similar strengths of inputs to each of the 74 regions (Hlinka & Coombes (2012), Forrester et al. (2020)).

#### 2.3.2 Model simulations

We used the Brain Dynamics Toolbox (BDT) for model simulations (Heitmann et al. (2018)). The Wilson-Cowan zero delays model is implemented in BDT, while we extended the BDT implementation of the zero delays model to implement the Wilson-Cowan constant delays and Wilson-Cowan distance-dependent delays models. We have made these implementations publicly available (Williams et al. (2021a)). We simulated the models for 630 seconds at 500 Hz with ODE45 (Bogacki & Shampine (1996)) for the Wilson-Cowan zero delays model, and ODE23a (Shampine & Thompson (2001)) for the constant delays and distance-dependent delays models. We limited local discretisation error by setting Absolute and Relative Tolerance of all solvers to 1 × 10^−6^ and 1 × 10^−3^ respectively. We set initial conditions of excitatory and inhibitory populations of the Wilson-Cowan oscillators by sampling a uniform random distribution between 0 and 1. Notably, we used dynamics of only the excitatory populations for further processing since pyramidal neurons in excitatory populations are the dominant contributors to the measured MEG (Lopes da Silva (2013)). These model dynamics represented the mean firing rate of pyramidal neurons in excitatory populations.

#### 2.3.3 Transforming simulated data to MEG source-space

We transformed simulated data to MEG source-space by forward-projecting the simulated data to MEG sensors and then inverse-projecting the simulated sensor-level data, with subject-specific forward and inverse operators respectively (Korhonen et al. (2014)). We used the same forward and inverse operators estimated to source-reconstruct the sensor-level MEG resting-state data (see Section 2.4.2). This procedure yielded subject-specific source-reconstructed simulated MEG datasets, of activity from 74 regions for 630 seconds.

#### 2.3.4 Band-pass filtering

We band-pass filtered the model dynamics in the alpha frequency band (8 - 12 Hz) since it is the dominant source of oscillatory power in MEG resting-state (Nakagawa et al. (2014)). We performed the filtering with Morlet wavelets of peak frequency = 9.83 Hz, width parameter = 5, yielding subject-specific narrowband datasets, from 74 regions for 630 seconds.

#### 2.3.5 Estimating network of phase-synchronization

For each subject-specific dataset, we used weighted Phase Lag Index (wPLI) (Vinck et al. (2011)) to estimate strength of phase-synchronization between each pair of brain regions.

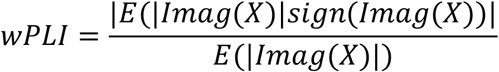

where *X* is the cross-spectrum between a pair of signals and *imag*(*X*) is its imaginary component. Crucially, wPLI is insensitive to the confounding influence of MEG field spread (Vinck et al. (2011), Palva et al. (2018)). We removed the effect of transient dynamics on the phase-synchronization estimates by discarding the first 30 seconds of each dataset. The phase-synchronization estimation produced 74 × 74 subject-specific matrices, which we averaged to obtain the model-generated network of phase-synchronization.

### 2.4 Obtaining MEG-derived network of phase-synchronization

Below, we describe the details of acquiring, preprocessing, source-reconstructing sensor-level MEG resting-state data and estimating the MEG-derived network of phase-synchronization.

#### 2.4.1 Data acquisition & pre-processing

We recorded MEG resting-state data from 46 healthy subjects (19 female, age 30.7 ± 9.5 years) for 600 seconds at a sampling rate of 1000 Hz. During the recordings, we instructed subjects to focus on a cross on the screen in front of them during the recordings. We performed the recordings at Meilahti hospital, Helsinki with a 306-channel MEG system (204 planar gradiometers and 102 magnetometers, MEGIN Oy). For each subject, we collected T1-weighted anatomical MRI scans at a resolution of 1 × 1 × 1 mm with a 1.5T MRI scanner (Siemens, Germany). The Ethics Committee of Helsinki University Central Hospital approved the study, and we performed the study according to the Declaration of Helsinki. We obtained written informed consent from each participant. We used MaxFilter (Taulu & Hari (2009)) to suppress extra-cranial noise and co-localise the sensor-space recordings, and Independent Component Analysis (ICA) (Oostenveld et al. (2011)), to exclude components identified as ocular or cardiac artefacts.

#### 2.4.2 Estimating MEG-derived network of phase-synchronization

We transformed preprocessed, sensor-level MEG data time series to brain regions, then band-pass filtered these data and estimated networks of phase-synchronization. We used MNE software to generate subject-specific inverse operators (Hämäläinen & Ilmoniemi (1994)) to transform sensor-level data to the level of cortical parcels or brain regions (Palva et al. (2010), Zhigalov et al. (2017), Lobier et al. (2018), Hirvonen et al. (2018)). We applied fidelity weighting (Korhonen et al. (2014)) on the inverse operators to reduce the influence of MEG field spread. We also used MNE to generate subject-specific forward operators, including subject-specific head conductivity models, cortically constrained source models with 5000-7500 sources per hemisphere, and to co-localise MEG to MRI. We used FreeSurfer (http://surfer.nmr.mgh.harvard.edu/) for volumetric segmentation of MRI data, surface reconstruction, flattening, cortical parcellation and neuroanatomical labelling of each source to a region in the Destrieux brain atlas. For each subject, we averaged the time series of sources within a region to obtain the activity time-course for that region. We then downsampled each subject-specific dataset to 500 Hz and retained the time courses of the 74 left-hemispheric regions for further processing. We used an identical procedure as applied to the simulated data, to perform band-pass filtering (see Section 2.3.4) and to estimate subject-specific 74 × 74 MEG-derived matrices of phase-synchronization (see Section 2.3.5).

### 2.5 Splitting MEG-derived networks into training and testing datasets

We generated two subsets of the original 46 subject-specific phase-synchronization matrices, to estimate free model parameters and to evaluate model performance respectively. The first subset or training dataset contained 23 unique, randomly selected subject-specific matrices. We averaged the matrices to obtain a group-level matrix, which we used as the target MEG-derived network to estimate model parameters. The held-out or testing dataset contained the 23 subject-specific matrices not used in the first subset. We averaged these matrices to obtain a group-level matrix, which we used as the target MEG-derived network to determine correspondence between model-generated and MEG-derived networks.

### 2.6 Determining optimal values of model parameters

We selected optimal model parameter values by performing a grid search for a range of parameter values. We quantified correspondence between model-generated and MEG-derived networks, *i*.*e*., model performance, as RMSE and Pearson Correlation between the upper-triangular elements of the model-generated and the training set MEG-derived matrices of phase-synchronization (see Section 2.5). We chose optimal parameter values as those simultaneously yielding low RMSE and high Correlation. For the zero delays, constant delays and distance-dependent delays models, we used the parameter range of *k* = 0.5 to 5, in intervals of 0.5, for the scalar multiplier over the structural connectome. For the constant delays model, we had a delay parameter *T*, with range from 2 ms to 20 ms, in intervals of 2 ms. For the distance-dependent delays model, we had a conduction velocity parameter *ν*, with range from 3 m/s to 30 m/s, in intervals of 3 m/s. We ran simulations for 315 seconds for each parameter combination and discarded the first 30 seconds to remove transient model dynamics, before estimating the model-generated network of phase-synchronization.

### 2.7 Evaluating statistical significance of model performance

We evaluated statistical significance of model performance by comparing against permutation-based null distributions of the performance measures. We quantified model performance by RMSE and Correlation between upper-triangular elements of the mean model-generated matrix of phase-synchronization and the held-out testing set MEG-derived matrix of phase-synchronization (see Section 2.5). We estimated the mean model-generated matrix of phase-synchronization by averaging the 20 model-generated matrices from different starting conditions. We determined statistical significance of the RMSE and Correlation values by *z*-scoring them against 100 corresponding values estimated after randomly permuting, without replacement, the mean model-generated matrix. We tested the hypotheses that RMSE is lower than chance and that Correlation is higher than chance by calculating the one-tailed *p*-value of the *z*-scores, with *p* < 0.001 considered statistically significant.

### 2.8 Sensitivity of model performance to choice of model parameter values

We assessed the sensitivity of the model’s performance to small changes in parameter values. To do this, we determined RMSE and Correlation values between the MEG-derived matrix and model-generated matrices generated from simulating jittered models. To introduce jitter, each non-zero parameter of the model was varied between -10% and 10% of its original value, with 2% increments, while keeping all other parameters at their original values. Then, we *z*-scored the RMSE and Correlation values against the 20 RMSE and Correlation values from the original model. *z* values higher than 3 were considered to reflect values of parameters at which the model performance was sensitive.

### 2.9 Determining features of each model component contributing to model performance

For the model producing networks of phase-synchronization corresponding most closely to the MEG-derived networks of phase-synchronization, we determined specific features of each model component, *e*.*g*., pattern of connections in the structural connectome, contributing to the observed correspondence. To do this, we compared the correspondence between model-generated and MEG-derived networks obtained with the original model, against the correspondence obtained with 100 examples of a null model, each containing a randomised version of a specific model component, *e*.*g*., randomly connected structural connectome, while all other aspects of the model remained identical to the original model.

Each of the Wilson-Cowan zero delays, constant delays and distance-dependent delays models, postulate that the topological organisation of the structural connectome, *i*.*e*., its pattern of connections, contributes to the correspondence between model-generated and MEG-derived networks. To verify this, we created 100 examples of a null model containing degree-preserved but randomised versions (Maslow & Sneppen (2002)) of the original binary structural connectome, while all other aspects were identical to the original model. For each example of this null model, the randomised structural connectome had the same number of connections to each brain region as the original structural connectome, but the pattern of connections between regions was different to the original structural connectome. By comparing the correspondence between model-generated and MEG-derived networks of the original model against the correspondences obtained with the null models, we tested the hypothesis that the pattern of structural connections contributed to the observed correspondence, over and above the number of connections, *i*.*e*., degree, to each brain region.

Each of the Wilson-Cowan zero delays, constant delays and distance-dependent delays models postulate that the Wilson-Cowan oscillatory dynamics contribute to the correspondence between model-generated and MEG-derived networks. To verify this, we created 100 examples of a null model where the dynamics of each brain region were described by a uniform random distribution whose mean (0.2) and range (0.07 to 0.36) were matched to those of original Wilson-Cowan dynamics, while the original structural connectome specified the interaction between regions. The model is specified as:

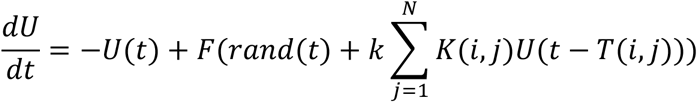

where 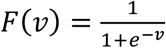 is a sigmoid function, *U*(*t*) is the mean firing rate, *rand*(*t*) is a sample from a uniform random distribution, *k* is a scalar multiplier over the coupling matrix *K*, which specifies links between *N* oscillators and *T*(*i, j*) is a delay matrix containing zeros for all elements, for the Wilson-Cowan zero delays model, a constant non-zero delay for all elements, for the Wilson-Cowan constant delays model, and connection-specific delays for each element, for the Wilson-Cowan distance-dependent delays model. We estimated the parameter *k* using the same procedure as for the original model (see Section 2.6). We set Relative Tolerance of the simulation to 0.1, to ensure completion in a reasonable time. By comparing the correspondence between model-generated and MEG-derived networks of the original model against the correspondences obtained with the null models, we tested the hypothesis that the Wilson-Cowan oscillatory dynamics contributed to the correspondence between model-generated and MEG-derived networks of phase-synchronization, over and above random dynamics with same mean and range as the Wilson-Cowan oscillatory dynamics.

The Wilson-Cowan distance-dependent delays model postulated that the connection-specific delays contributed to the correspondence between the model-generated and MEG-derived networks. To verify this, we created 100 examples of a null model by randomly permuting delays (without replacement) from the original model, while all other aspects, including the structural connectome and Wilson-Cowan oscillatory dynamics, were identical to the original model. By comparing the correspondence between model-generated and MEG-derived networks of the original model against the correspondences obtained with the null models, we tested the hypothesis that the connection-specific delays contributed to the observed correspondence, over and above merely the set of conduction delays used. Since the Wilson-Cowan zero delays and Wilson-Cowan constant delays models had the same delays for each connection, determining the contribution of connection-specific delays to the correspondence between model-generated and MEG-derived networks was not relevant.

Correspondence between the model-generated and MEG-derived networks, for both the original model and null models, were quantified by RMSE and Correlation between the upper-triangular elements of model-generated and the testing set MEG-derived matrices of phase-synchronization (see Section 2.5). To remove outliers from the null distribution, we excluded cases for which RMSE values lay more than 5SD from the mean. We then estimated the *z*-score of the original model’s RMSE and Correlation against RMSE and Correlation of the null models. We tested the hypotheses that mean RMSE for the original model is lower than RMSE for the null models and mean Correlation for the original model is higher than Correlation for the null models by calculating one-tailed p-values of the respective *z*-scores, with *p* < 0.001 considered a statistically significant contribution.

### 2.10 Comparing model performance against performance of Kuramoto oscillator model

For the model producing networks of phase-synchronization corresponding most closely to those observed in MEG resting-state, we compared its observed correspondence against the correspondence between model-generated and MEG-derived networks of an equivalent Kuramoto oscillator model (Kuramoto (1984)). The Kuramoto oscillator model has been used to model oscillatory phenomena in systems neuroscience (Breakspear et al. (2010), Cabral et al. (2014)). The phase dynamics of individual regions were described by the classic Kuramoto model, where we used the same coupling matrix *K* as in the Wilson-Cowan models, to specify links between regions. The sine of instantaneous phases was computed to convert the phase dynamics to time series of oscillatory activity. We set the scalar multiplier *k* over the coupling matrix with the same training procedure used for the Wilson-Cowan zero delays model (Section 2.6), while we set the natural frequencies of the Kuramoto oscillators so as to produce oscillatory dynamics around 10 Hz. To do this, we set the natural frequencies as independent samples from a Gaussian distribution of mean = 6.2 × *10*^−*2*^ and standard deviation = 6.2 × *10*^−*3*^. We simulated the model, then applied the same processing as to the Wilson-Cowan model dynamics (see Section 2.3) to furnish the Kuramoto model-generated matrix of phase-synchronization, which we then matched to the MEG-derived matrix with RMSE and Correlation. We used independent samples *t*-tests to compare RMSE and Correlation values of the chosen model against RMSE and Correlation values of the Kuramoto model, with two-tailed *p* < 0.001 considered statistically significant.

## 3. Results

### 3.1 Determining optimal values of model parameters

We estimated the optimal values of free parameters, for the Wilson-Cowan zero delays, constant delays and distance-dependent delays models. We identified the optimal parameter values as those that yielded model-generated matrices of phase-synchronization with the strongest correspondence to the training set MEG-derived matrices of phase-synchronization. For each of the models, we compared model-generated and MEG-derived matrices of phase-synchronization using RMSE and Pearson Correlation, choosing parameter values yielding low RMSE and high Correlation (Figure 2). For the Wilson-Cowan zero delays model, the optimal parameter value is *k* = 2, yielding RMSE = 0.04 and Correlation = 0.45, *k* being the scalar multiplier over the structural connectome. For the Wilson-Cowan constant delays model, the optimal combination of parameter values is *k* = 2 and *T* = 2 ms, yielding RMSE = 0.04 and Correlation = 0.33, with *T* being the constant delay between brain regions. For the Wilson-Cowan distance-dependent delays model, the optimal combination of parameter values is *k* = 1.5 and *ν* = 12 m/s, yielding RMSE = 0.04 and Correlation = 0.38, with *ν* being the assumed transmission velocity of neuronal activity across structural connections. We used these values to run fresh simulations of the Wilson-Cowan zero delays, constant delays and distance-dependent delays models, and determined correspondences of the respective model-generated matrices of phase-synchronization to the testing set MEG-derived matrix of phase-synchronization, which we estimated as the average of 23 held out subject-level matrices.

**Figure 2.**
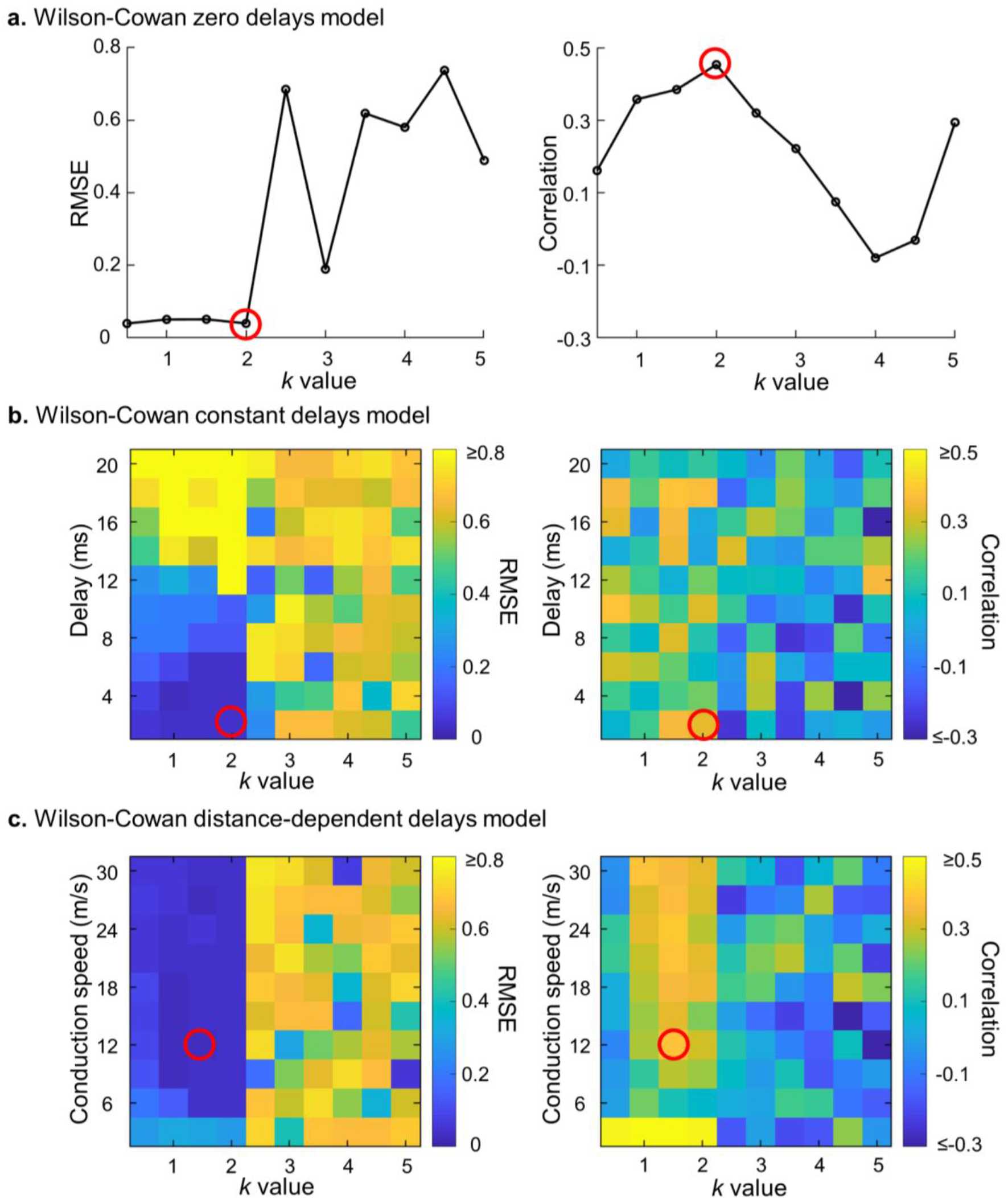
Grid search for optimal parameter values of each model. a. RMSE and Correlation values for a range of *k* values, for Wilson-Cowan zero delays model b. RMSE and Correlation values for every combination of a range of *k*-values and a range of delay values, for Wilson-Cowan constant delays model c. RMSE and Correlation values for every combination of a range of *k*-values and range of conduction speed values, for Wilson-Cowan distance-dependent delays model. *k* is the scalar multiplier over the structural connectome.

### 3.2 Model comparison

We compared the Wilson-Cowan zero delays, Wilson-Cowan constant delays and Wilson-Cowan distance-dependent delays models. To do this, we compared the correspondences between their respective model-generated networks of phase-synchronization, and the testing set MEG-derived network of phase-synchronization. We quantified correspondence with RMSE and Correlation between upper triangular elements of the model-generated matrices from 20 simulations of each model and the testing set MEG-derived matrix. We used independent samples *t*-tests to compare performances between models and Cohen’s *d* to measure effect sizes. RMSE values from the constant delays model (mean = 0.041) are higher than RMSE values from both the zero delays (mean = 0.039) (two-tailed *p* = 1.1 × 10^−8^, Cohen’s *d* = 2.3) and distance-dependent delays models (mean = 0.039) (two-tailed *p* = 7.4 × 10^−5^, Cohen’s *d* = 1.4) (Figure 3, left panel). In contrast, there is no difference between RMSE values from the zero delays and distance-dependent delays models (two-tailed *p* > 0.001, Cohen’s *d* = 0.37). Similarly, Correlation values from the constant delays model (mean = 0.37) are lower than Correlation values from the zero delays (mean = 0.47) (two-tailed *p* = 4.7 × 10^−10^, Cohen’s *d* = 2.62) and distance-dependent delays models (mean = 0.46) (two-tailed *p* = 1.6 × 10^−10^, Cohen’s *d* = 2.74) (Figure 3, right panel), while there is no difference in Correlation values from the zero delays and distance-dependent delays models (two-tailed *p* > 0.001, Cohen’s *d* = 0.12). These results suggest that the presence of delays in the Wilson-Cowan distance-dependent delays model does not improve the correspondence between model-generated and MEG-derived networks, compared to the correspondence obtained with the Wilson-Cowan zero delays model. Hence, we chose the more parsimonious model, *i*.*e*. the Wilson-Cowan zero delays model, for further investigation.

**Figure 3.**
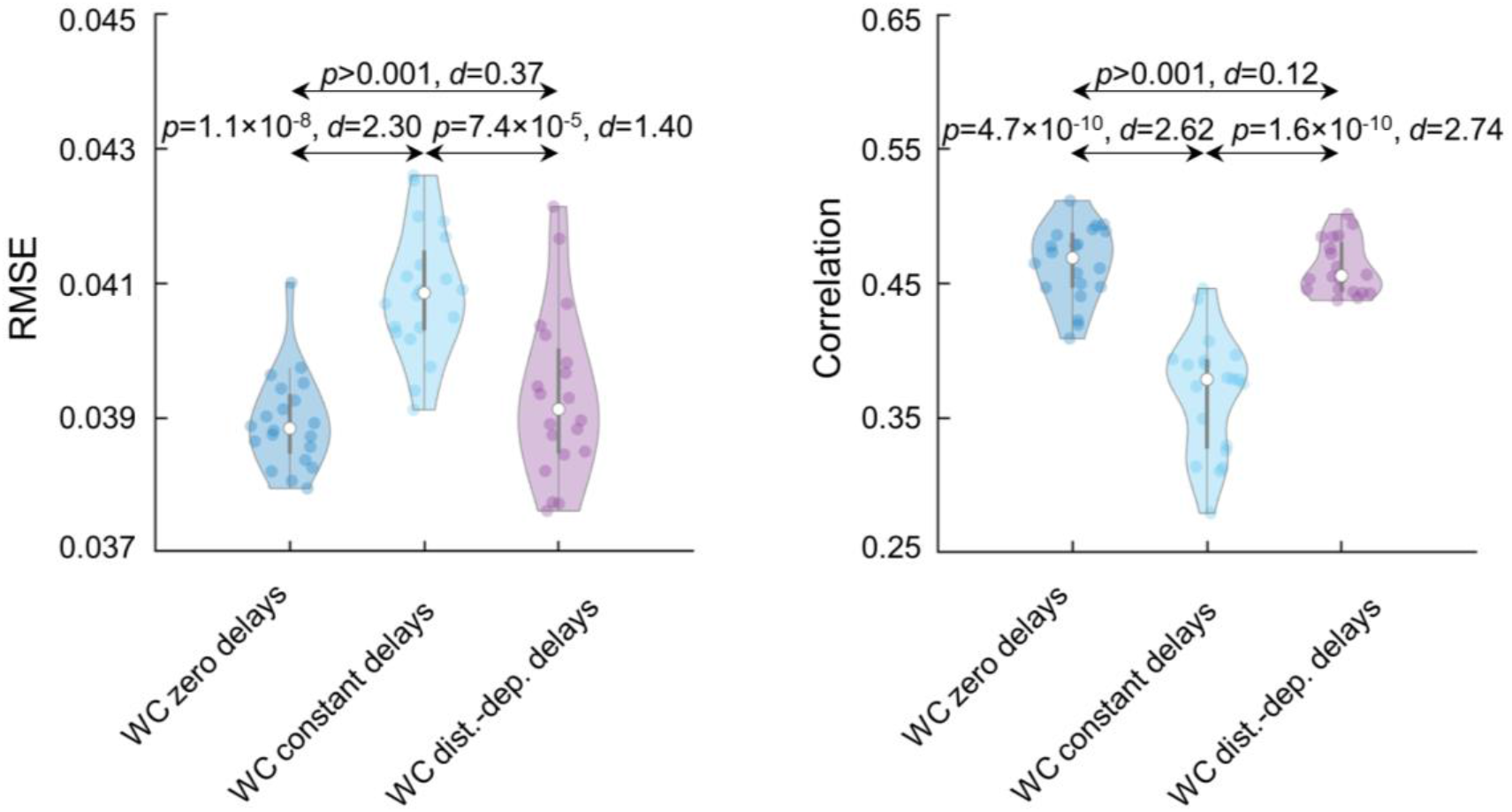
Including conduction delays does not improve model performance. Violin plots of RMSE values for the Wilson-Cowan zero delays (mean = 0.039), constant delays (mean = 0.041) and distance-dependent delays models (mean = 0.039) (left panel). Violin plots of Correlation values for the Wilson-Cowan zero delays (mean = 0.47), constant delays (mean = 0.37) and distance-dependent delays models (mean = 0.46) (right panel).

### 3.3. Close correspondence between model-generated and MEG-derived network of phase-synchronization

We observed that the source-reconstructed dynamics of the Wilson-Cowan zero delays model exhibited intermittent dynamics (Figure 4a), also observed in MEG resting-state data (Shriki et al. (2013)). Further, the frequency spectrum of the dynamics displayed peaks in delta (1 - 4 Hz), alpha (8 - 12 Hz) and beta (12 - 30 Hz) frequency bands (Figure 4b), which have also been observed in MEG resting-state data (Mahjoory et al. (2020), Lopes da Silva (2013)).

**Figure 4.**
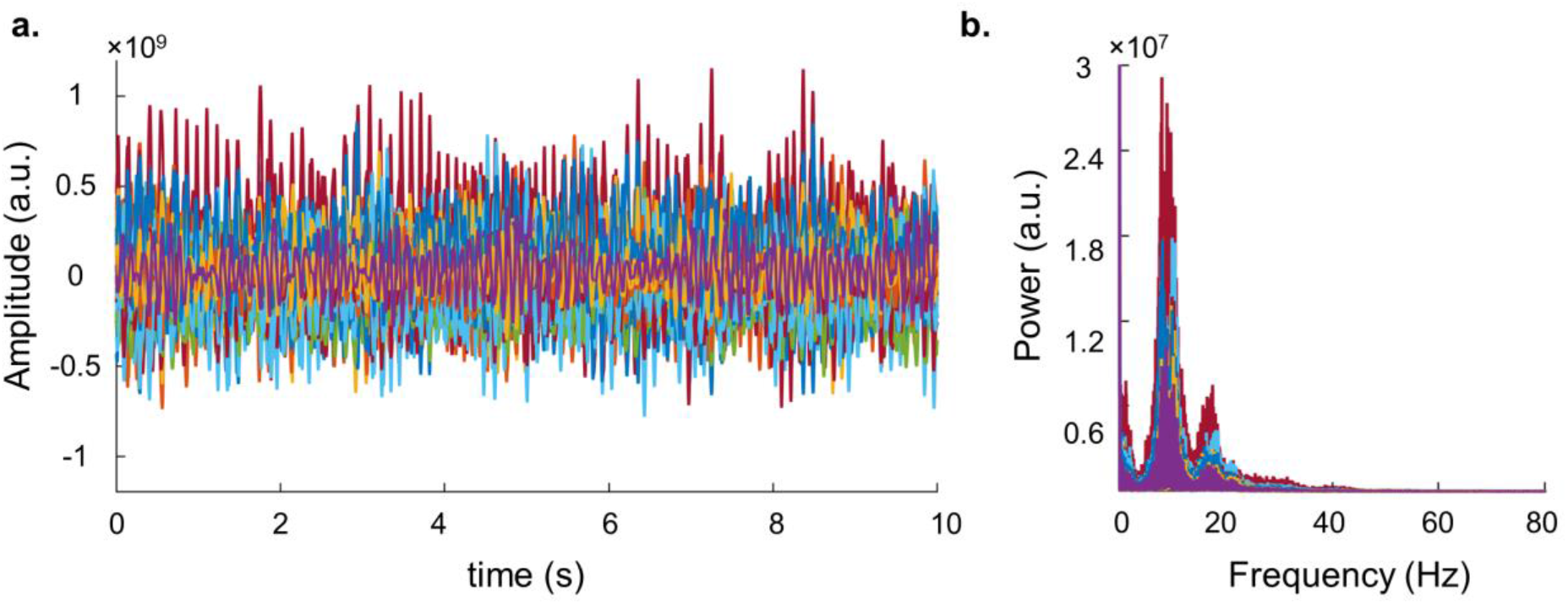
Intermittent dynamics and peaks in the frequency spectrum of dynamics of Wilson-Cowan zero delays model are similar to those observed in MEG resting-state. a. 10-second time course of model dynamics from all brain regions for example subject b. Frequency spectrum of model dynamics for 10-second segment of model dynamics from all brain regions, for example subject. Different colours indicate time course/frequency spectrum of different brain regions. Same colour in panels a, b belongs to the same brain region.

We compared the levels of phase-synchronization within the source-reconstructed dynamics of the Wilson-Cowan zero delays model to those observed in source-reconstructed MEG resting-state data. To do this, we compared the mean of the Kuramoto order parameter (Breakspear et al. (2010)) of the source-reconstructed model dynamics and source-reconstructed MEG resting-state data, across participants in the held-out testing set. We observed mean of the Kuramoto order parameter of the model dynamics (Figure 4a) (mean = 8 × 10^−2^, standard deviation = 9 × 10^−3^) to be close to the mean of the Kuramoto order parameter of the source-reconstructed MEG resting-state data (mean = 8 × 10^−2^, standard deviation = 1 × 10^−2^). Hence, the source-reconstructed dynamics of the Wilson-Cowan zero delays model display similar levels of phase-synchronization as those observed in source-reconstructed MEG resting-state.

We next assessed the correspondence between the mean model-generated and MEG-derived networks of phase-synchronization. The mean model-generated network was the average of 20 model-generated networks, each estimated from simulated model data from different initial conditions. We compared the mean model-generated network of phase-synchronization to the testing set MEG-derived network. Specifically, we compared the distributions of mean model-generated and MEG-derived phase-synchronization values, and compared the RMSE and Correlation between the mean model-generated and MEG-derived matrices against their corresponding permutation-based null distributions. The distributions of mean model-generated and MEG-derived phase-synchronization values are similar, with central tendency (median = 0.09) and dispersion (median absolute deviation = 0.02) of mean model-generated network values close to central tendency (median = 0.07) and dispersion (median absolute deviation = 0.02) of MEG-derived network values (Figure 5a). RMSE = 0.04 between mean model-generated and MEG-derived matrices is lower than chance (*z* = -29.9, one-tailed *p* = 6.7 × 10^−197^), and Correlation = 0.49 between mean model-generated and MEG-derived networks is higher than chance (*z* = 25.4, one-tailed *p* = 0) (Figure 5b,c). Further, most top 5 percentile strongest connections lie within and between parietal, temporal and occipital regions, for both mean model-generated and MEG-derived networks of phase-synchronization (Figure 5d). Taken together, the results demonstrate a close correspondence between the mean model-generated and MEG-derived networks of phase-synchronization.

**Figure 5.**
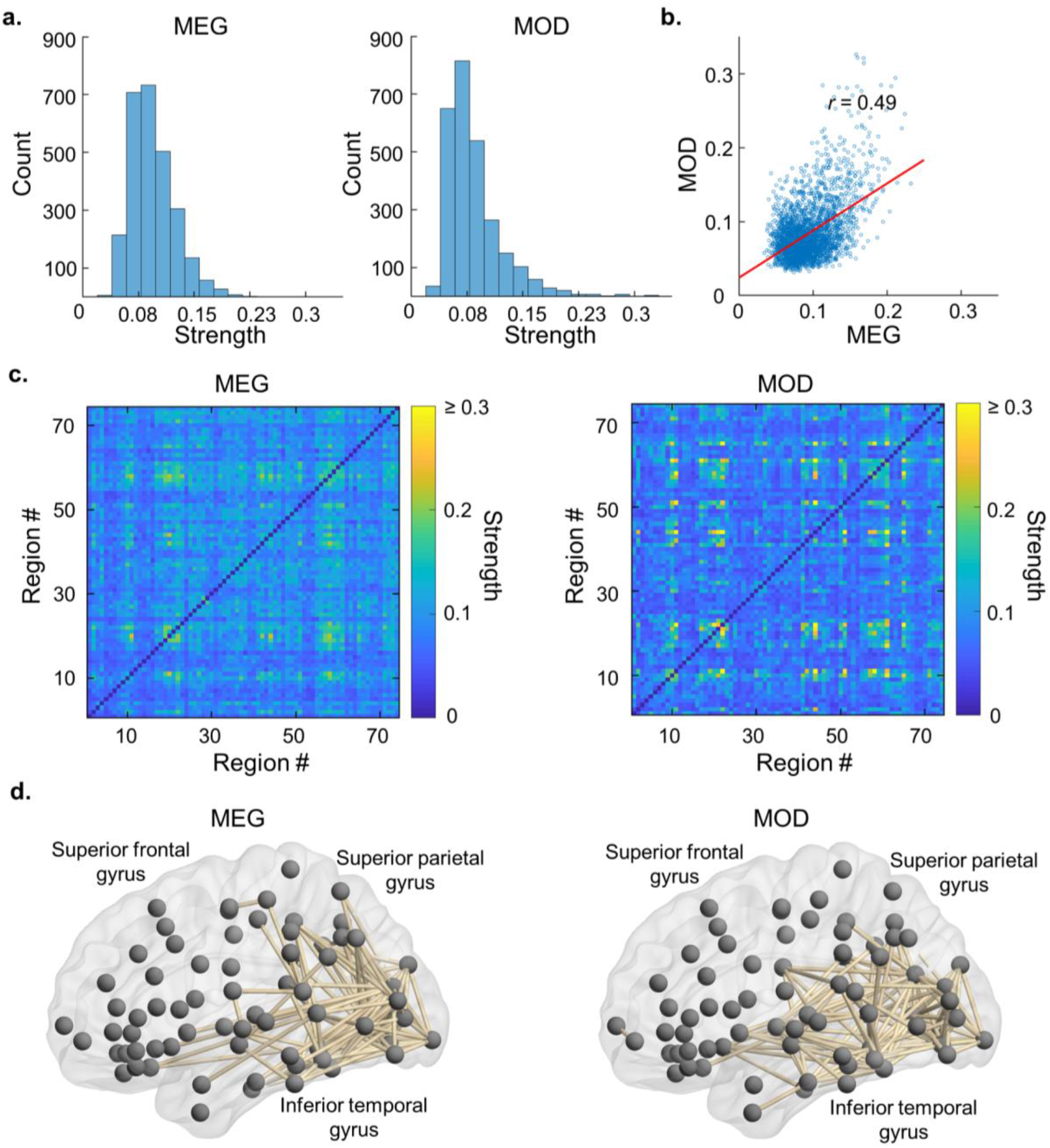
Close correspondence between mean model-generated network and MEG-derived network of phase-synchronization. a. Histogram of connection strengths for mean model-generated and MEG-derived networks of phase-synchronization b. Scatter plot of connection strengths for MEG-derived network of phase-synchronization against mean model-generated network of phase-synchronization c. Matrices of connection strengths between every pair of brain regions for mean model-generated and MEG-derived network of phase-synchronization. d. Brain network visualisation of top 5 percentile strongest connections for mean model-generated and MEG-derived networks of phase-synchronization. MOD = model. Brain networks were visualised with BrainNet Viewer (http://www.nitrc.org/projects/bnv/) (Xia et al. (2013)).

We assessed the sensitivity of the Wilson-Cowan zero delays model, to changes in the values of model parameters. To do this, we performed a sensitivity analysis wherein we estimated change in model performance to small changes in the values of model parameters. We varied the value of each non-zero parameter from - 10% to 10% of its original value, while keeping all other parameters at their original value. The RMSE and Correlation from simulating these jittered models were *z*-scored against the 20 RMSE and Correlation values from simulating the original model, to estimate the change in model performance. We found the model performance to be robust to changes in values of *k* and *w*_*ii*_ parameters (*z* values < 3), *i*.*e*. the scalar multiplier over the structural connectome and weight of self-inhibitory connections of Wilson-Cowan oscillators respectively. However, the model performance is sensitive to changes in the values of all other model parameters (*z* values > 3) (Figure S1). Hence, we observe that the Wilson-Cowan zero delays model is sensitive to changes in the values of its model parameters.

We have shared the mean model-generated network and networks from each of 20 simulations of the Wilson-Cowan zero delays model, through an open dataset (Williams et al. (2021b)).

### 3.4 Determining features of each model component contributing to model performance

The Wilson-Cowan zero delays model represents the hypothesis that the Wilson-Cowan oscillatory dynamics interact with the pattern of structural connections to produce networks of phase-synchronization corresponding to MEG-derived networks of phase-synchronization. Hence, it postulates that both the pattern of structural connections and Wilson-Cowan oscillatory dynamics contribute to the observed correspondence.

We tested the hypothesis that the pattern of structural connections contributes to the correspondence between model-generated and MEG-derived networks. To do this, we compared correspondence obtained with the original model against correspondence obtained with null models containing degree-preserved randomised versions of the structural connectome, all other aspects of the null model being identical to the original model. The structural connectome from each of the null models had the same number of connections to each brain region as the original structural connectome, but the pattern of connections between regions was different from that of the original structural connectome. We used RMSE and Correlation between the model-generated and testing set MEG-derived matrices to quantify correspondence. Mean RMSE with the original model of 0.04 was lower (*z* = -3.7, one-tailed *p* = 1 × 10^−4^) and mean Correlation of 0.47 was higher (*z* = 8.4, one-tailed *p* = 0) than the set of RMSE and Correlation values yielded by null models containing degree-preserved randomised versions of the original structural connectome (Figure 6a). These results confirm that the pattern of connections in the structural connectome contributes to the observed correspondence between model-generated and MEG-derived networks, over and above the number of connections, *i*.*e*. degree, to each brain region.

**Figure 6.**
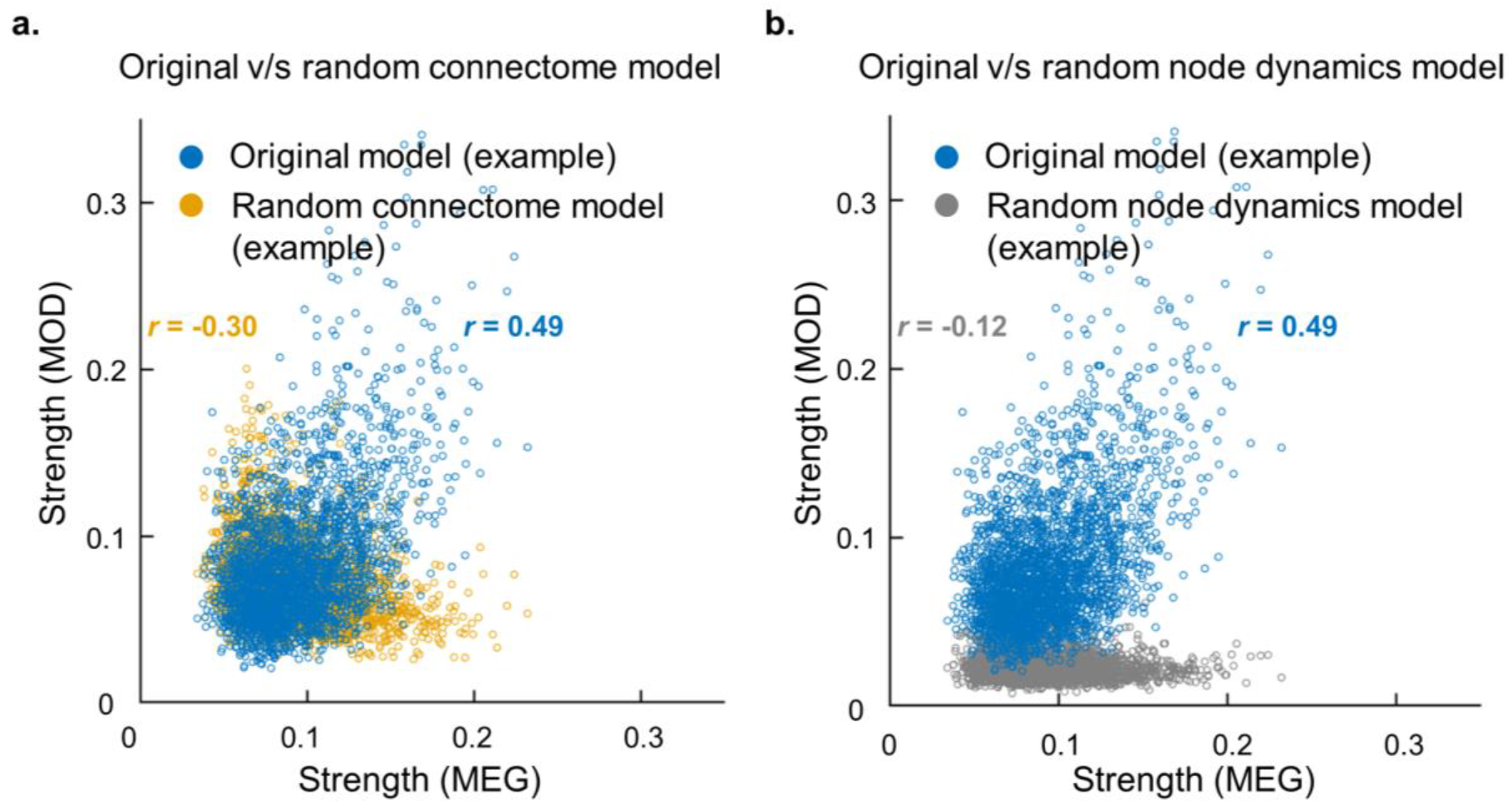
The pattern of connections in the structural connectome and dynamics of Wilson-Cowan oscillators, contribute to correspondence between model-generated and M EG-derived networks of phase-synchronization. a. Scatter plot of connection strengths from MEG-derived network and example model-generated network (blue) and of connection strengths from MEG-derived network and example null model-generated network, wherein structural connectome is randomised (yellow). b. Scatter plot of connection strengths from MEG-derived network and example model-generated network (blue) and scatter plot of connection strengths from MEG-derived network and null model-generated network, wherein node dynamics are randomised (dark gray). MOD = model.

Given the contribution of the pattern of connections of the structural connectome to the observed correspondence between model-generated and MEG-derived networks, we investigated if the model-generated network is merely a recapitulation of the structural connectome. To do this, we compared Correlation between the model-generated matrix and the testing set MEG-derived matrix, against Correlations between 100 bootstrapped versions of the structural connectome matrix and the testing set MEG-derived matrix. Mean Correlation of 0.47 between the model-generated and MEG-derived matrices is higher (*z* = 181.5, one-tailed *p* = 0) than Correlations between bootstrapped versions of the structural connectome matrix and the MEG-derived matrix (Figure S2). These results demonstrate that the model does not merely recapitulate the structural connectome at the level of its dynamics, but that it operates on the structural connectome in a non-trivial way to produce the observed MEG-derived matrix of phase-synchronization.

We tested the hypothesis that the Wilson-Cowan oscillatory dynamics contributes to the comparison between model-generated and MEG-derived networks of phase-synchronization. To do this, we compared correspondence obtained with the original model against correspondence obtained with null models wherein the dynamics of each brain region were described by random dynamics interacting through the structural connectome used in the original model. The mean and range of random dynamics were matched to those of the Wilson-Cowan oscillators. For the random node dynamics model also, we estimated the model parameter *k*, the scalar multiplier over the structural connectome, with the same procedure as for the Wilson-Cowan zero delays model (Section 2.6). We used RMSE and Correlation between the model-generated and testing set MEG-derived matrices to quantify correspondence. Note that Correlation is insensitive to scale of the compared phase-synchronization strengths, hence it would yield high values for similar patterns of model-generated and MEG-derived phase-synchronization even if the random node dynamics model produced weaker phase-synchronization strengths than those observed in the MEG-derived networks. Just as for the Wilson-Cowan zero delays model, the optimal value of *k* for the random node dynamics model was identified as 2. Mean RMSE with the original model of 0.04 was lower (*z* = -40.8, one-tailed *p* = 0) and mean Correlation of 0.47 was higher (*z* = 4.9, one-tailed *p* = 5 × 10^−7^) than the set of RMSE and Correlation values yielded by null models containing random node dynamics instead of Wilson-Cowan oscillatory dynamics (Figure 6b). These results confirm that the Wilson-Cowan oscillatory dynamics contributes to the observed correspondence between model-generated and MEG-derived networks, over and above dynamics with the same mean and range.

Given the contribution of Wilson-Cowan oscillator dynamics to the correspondence between model-generated and MEG-derived networks, we investigated if the mere presence of oscillatory rather than random node dynamics would produce the observed correspondence. To do this, we compared correspondence obtained with the Wilson-Cowan zero delays model against correspondence obtained with an equivalent Kuramoto oscillator model, which also produces oscillatory dynamics. No statistically significant difference in the correspondences with the Wilson-Cowan zero delays model and Kuramoto oscillator model would imply the mere presence of oscillatory node dynamics produces a correspondence between model-generated and MEG-derived networks. We quantified correspondence using the RMSE and Correlation between the respective model-generated and MEG-derived matrices of phase-synchronization. We identified *k* = 0.5 as the optimal value of the scalar multiplier over the structural connectome for the Kuramoto oscillator model. Mean RMSE of 0.04 with the Wilson-zero delays model was lower than mean RMSE of 0.53 with the Kuramoto oscillator model (two-tailed *p* = 8.4 × 10^−59^). Similarly, mean Correlation of 0.47 with the Wilson-Cowan zero delays model is higher than mean Correlation of 0.13 with the Kuramoto oscillator model (two-tailed *p* = 2.2 × 10^−37^) (Figure S3). These results demonstrate the mere presence of oscillatory node dynamics does not produce the observed correspondence between model-generated and MEG-derived networks. Rather, it is the oscillatory dynamics of the Wilson-Cowan oscillators resulting from the interaction between excitatory and inhibitory neuronal populations, which produces the observed correspondence.

### 3.5 Robustness of results

Finally, we investigated the robustness of the obtained correspondence between model-generated and MEG-derived networks, to choices made for the simulations and analyses. To do this, we compared correspondence obtained with different solvers and with different relative tolerance values for the model simulations. In addition, we determined if changing the modeled hemisphere from left to right would qualitatively change the correspondence between the model-generated and MEG-derived networks. We quantified correspondence by the RMSE and Correlation between the model-generated and MEG-derived matrices of phase-synchronization. We used the ODE45 solver in the original simulations, but RMSE and Correlation values are not different when ODE23 and ODE113 solvers were used (two-tailed *p* > 0.001 for each) (Figure 7a). Similarly, we used relative tolerance of 10^−3^ for the original simulations, which yielded RMSE and Correlation values neither different to those obtained with lower relative tolerance values of 10^−4^ and 10^−5^ nor a higher relative tolerance of 10^−2^ (two-tailed *p* > 0.001 for each) (Figure 7b). Changing the modeled hemisphere from left to right also yielded a moderate correspondence between the model-generated and MEG-derived network (Correlation = 0.33) (Figure S4). Hence, the correspondence between model-generated and MEG-derived networks of phase-synchronization, were robust to choices made for the simulations and analyses.

**Figure 7.**
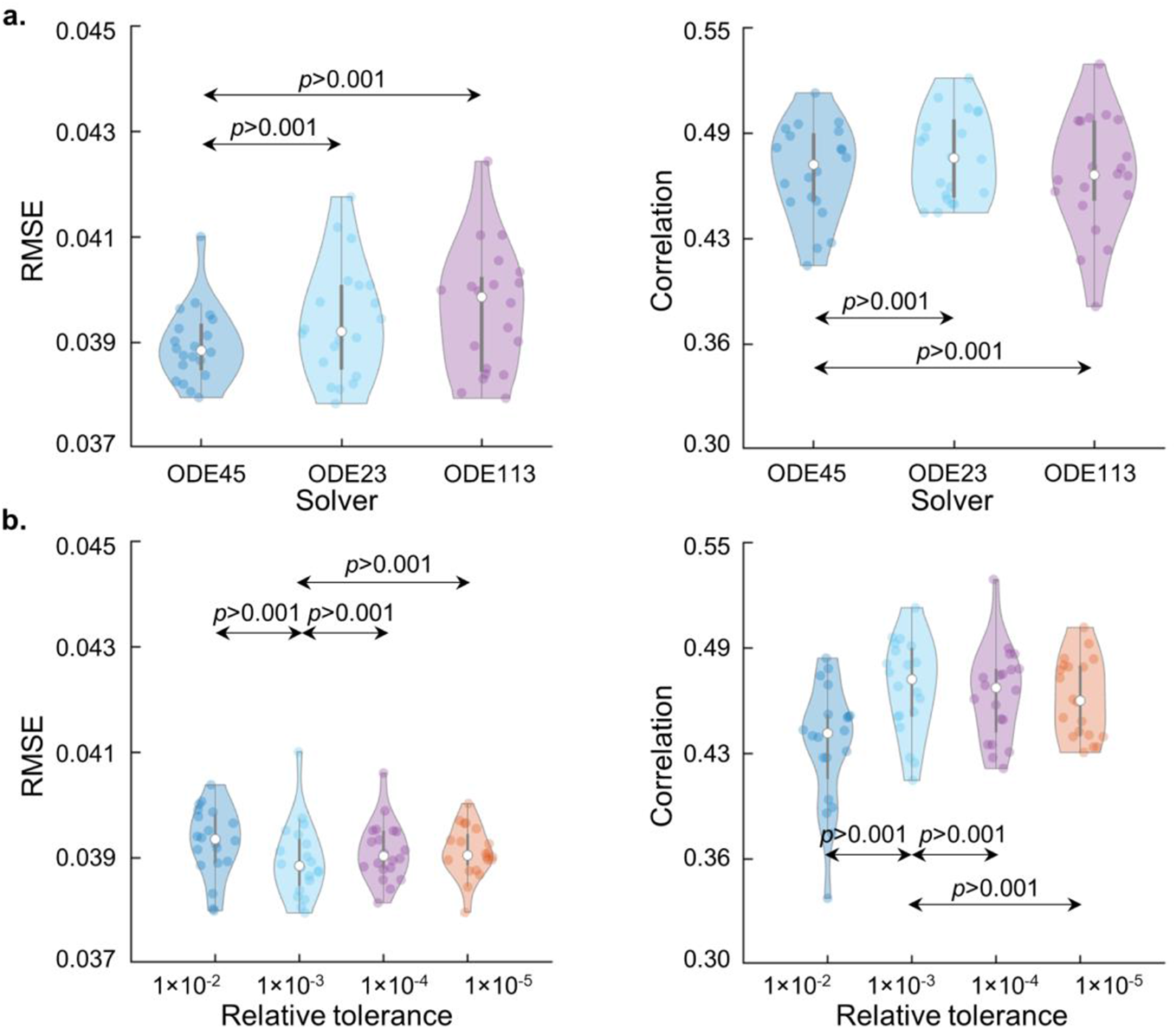
Results are robust to changes in solver and relative tolerance of solution. a. Boxplot of RMSE and Correlation with different solvers. Solver used in the original model was ODE45. b. Boxplot of RMSE and Correlation at different relative tolerances of solution. Relative tolerance used in the original model was 10^−3^.

## 4. Discussion

Large-scale networks of phase-synchronization are widely observed with EEG or MEG during rest and task. BNMs are used to produce model-generated networks corresponding to MEG-derived networks of phase-synchronization, but the respective roles of conduction delays, the structural connectome and dynamics of models of individual brain regions, in obtaining this correspondence remain unknown. In this paper, we investigated the roles of conduction delays, the structural connectome and dynamics of models of individual regions, in producing networks of phase-synchronization corresponding to those observed in MEG resting-state. We found no evidence that including conduction delays improves the correspondence between the model-generated and MEG-derived networks of phase-synchronization. Further, we demonstrated that both the pattern of structural connections and Wilson-Cowan oscillatory dynamics contribute to obtaining the observed correspondence between model-generated and MEG-derived networks of phase-synchronization.

The topological organisation of the structural connectome, *i*.*e*., its pattern of connections, contributes to the observed correspondence between the model-generated and MEG-derived network of phase-synchronization. Previous studies have used the structural connectome within BNMs to produce other phenomena, such as networks matching experimentally observed fMRI-derived networks (Deco et al. (2009)), MEG-derived networks of amplitude correlation (Tewarie et al. (2019)) and spatial distribution of oscillation amplitudes observed in MEG resting-state (Raj et al. (2019)). Studies have also used the structural connectome in models to produce networks matching MEG-derived networks of phase-synchronization, but these have been via abstract statistical or oscillator models (Finger et al. (2016)), graph theory measures (Wodeyar & Srinivasan (2019)) or formal series expansions (Meier et al. (2016)). A single study used the structural connectome with a biologically plausible model to produce networks of phase-synchronization matching those observed in the MEG resting-state (Abeysuriya et al. (2018)). Our study advances findings from these studies in the following ways: 1.) We used comparison against null models to identify the specific feature of the structural connectome, *i*.*e*., its topological organisation, rather than lower-level properties such as the degree of each brain region, contributing to the observed correspondence between model-generated and MEG-derived networks of phase-synchronization, and 2.) We demonstrated that the correspondence between the model-generated and MEG-derived networks of phase-synchronization is not explained merely by the correspondence between the structural connectome and the MEG-derived network of phase-synchronization. Consistent with computational modelling on the relationship between the structural connectome and networks of phase-synchronization (Forrester et al. (2020)), these results suggest a significant but nontrivial relationship between the structural connectome and the model-generated network of phase-synchronization.

The dynamics of Wilson-Cowan oscillators contribute to the observed correspondence between model-generated and MEG-derived networks of phase-synchronization. Previous modelling studies have studied the role of the structural connectome in producing observed functional networks, but the role of dynamics of brain regions in producing these networks has not received much attention. A recent computational modelling study (Forrester et al. (2020)) demonstrated the influence of node dynamics on the relationship between the structural connectome and model-generated networks of phase-synchronization. The study demonstrated that oscillatory node dynamics are accompanied by a nontrivial relationship between the structural connectome and model-generated networks of phase-synchronization. Our study advances previous work in the following ways: 1.) While previous work has related oscillatory node dynamics to model-generated networks, this is the first work demonstrating the importance of oscillatory node dynamics in producing networks of phase-synchronization corresponding to those observed in experimental MEG resting-state. 2.) We illustrate the importance of specifically Wilson-Cowan oscillatory dynamics, by demonstrating that the Wilson-Cowan zero delays model produces networks with closer correspondence to MEG-derived networks of phase-synchronization than does an equivalent model of Kuramoto oscillators. This result is consistent with computational modelling demonstrating the influence of both amplitude and phase dynamics, as produced by Wilson-Cowan oscillators, on estimates of phase-synchronization (Daffertshofer & van Wijk (2011)). Kuramoto oscillators only yield descriptions of phase dynamics. Hence, the oscillatory dynamics resulting from the interaction between excitatory and inhibitory neuronal populations, as described by Wilson-Cowan oscillators, contributes to producing networks of phase-synchronization corresponding to those observed in MEG resting-state.

We find no evidence that including conduction delays improves the correspondence between model-generated and MEG-derived networks of phase-synchronization. Modelling studies investigating the role of delays in producing networks resembling fMRI-derived functional networks or MEG-derived networks of amplitude correlation have yielded differing results. Results of some studies suggest the importance of delays in producing fMRI-derived networks (Ghosh et al. (2008), Deco et al. (2009)) or MEG-derived networks of amplitude correlation (Cabral et al. (2014), Nakagawa et al. (2014)), while other modelling studies not including delays also produce networks resembling fMRI-derived (Honey et al. (2007)) or MEG-derived networks of amplitude correlation (Deco et al. (2017)). The role of conduction delays in producing networks resembling MEG-derived networks of phase-synchronization has not received much attention. A recent modelling study (Abeysuriya et al. (2018)) reported that, in the special case of a high mean activity level (mean activity = 0.3) of Wilson-Cowan oscillators imposed by synaptic plasticity mechanisms, conduction delays were not necessary to producing networks resembling MEG-derived networks of phase-synchronization. In our study, we advance previous work by demonstrating that, also in the more general case of no specific mean target activity imposed by synaptic plasticity mechanisms, a close correspondence between model-generated and MEG-derived networks of phase-synchronization can be achieved.

The absence of evidence in our study, for the role of delays in producing MEG-derived networks of phase-synchronization, might be either due to errors in the estimates of conduction delay or due to conduction delays not being mechanistically important in producing MEG-derived networks of phase-synchronization. If the latter were true, one would expect the Wilson-Cowan constant delays and Wilson-Cowan distance-dependent delays model to produce networks corresponding equally to the MEG-derived network of phase-synchronization. However, the Wilson-Cowan distance-dependent delays model produces networks correspondingly more closely to the MEG-derived network, than networks produced by the Wilson-Cowan constant delays model. In fact, the Wilson-Cowan distance-dependent delays model produces networks corresponding as closely to the MEG-derived network, as networks produced by the Wilson-Cowan zero delays model. Hence, we propose that improved accuracy in estimating conduction delays, by accounting for axonal diameter and myelination (Nakagawa et al. (2014), Waxman & Swadlow (1977)) in addition to distance, could yield improved correspondence between networks produced by the Wilson-Cowan distance-dependent delays model and the MEG-derived network of phase-synchronization. Specialised diffusion MRI sequences could yield estimates of myelination and axonal diameter (Drobnjak et al. (2016), Whittall et al. (1997)), but these estimates remain elusive.

The Wilson-Cowan zero delays model compares well to previously proposed BNMs producing networks of phase-synchronization observed in the MEG resting-state. A strength of a recently proposed BNM (Abeysuriya et al. (2018)) with distance-dependent conduction delays and inhibitory synaptic plasticity, was that it used the same set of parameter values to simultaneously produce a correspondence to both MEG-derived networks of amplitude correlation and MEG-derived networks of phase-synchronization. However, two notable aspects of the Wilson-Cowan zero delays model we propose are 1.) its parsimonious nature in not including conduction delays or synaptic plasticity mechanisms, while still producing networks corresponding closely (ρ = 0.49) to MEG-derived networks of phase-synchronization and 2.) its strengths of phase-synchronization between regions were of the same order of magnitude and closer to those observed in the MEG-derived network (RMSE = 0.04) compared to previously proposed BNMs. However, a direct comparison between our model and those used in previous studies is difficult due to differences in the brain parcellation atlases used, number of hemispheres considered, measures of phase-synchronization used, as well as processing details of MEG and model-generated data.

A limitation of the Wilson-Cowan zero delays model we propose is that while it does produce networks corresponding to MEG-derived networks of phase-synchronization, it does not simultaneously produce networks of amplitude correlation nor produce spatial distribution of oscillation amplitudes corresponding to those observed in MEG resting-state (results not shown). However, we did find these phenomena were produced by the model operating at very low levels of structural coupling between regions, *e*.*g*., *k* values of 10^−3^, where *k* is the scalar multiplier across the structural connectome. This was outside the range of structural coupling strengths investigated in our study. Future studies will use likelihood-free inference methods to estimate parameters from wider parameter ranges (Lintusaari et al. (2017), Lintusaari et al. (2018), Gutmann & Corander (2016), Cranmer et al. (2020)). Sensitivity analysis of our model revealed the model performance to be highly sensitive to values of model parameters. This could be due to the absence, in the model, of empirically observed homeostatic mechanisms to counteract the effect of changes in *e*.*g*., the mean activity level of the Wilson-Cowan oscillators.

Future work could extend the model to include empirically observed spatial gradients in synaptic excitation (Wang (2020)), which in turn produce empirically observed spatial gradients in peak frequency (Hadida et al. (2019), Mahjoory et al. (2020)). The model could also be extended to include the effects of noise (Faisal et al. (2008)), which has been suggested to play a role in generating dynamics of resting-state data from other methodologies (Deco et al. (2009)). These models, fit to MEG resting-state data, could be locally perturbed to emulate task-related dynamics (Tiesinga et al. (2010)). Such virtual experiments could be used to develop, test and refine hypotheses on biophysical substrates underlying observed networks of phase-synchronization in MEG task data.

## 5. Conclusion

In this study, we investigated the respective contributions of conduction delays, the structural connectome and dynamics of models of individual brain regions, in producing model-generated networks of phase-synchronization corresponding to those observed in MEG resting-state. Based on our investigations, we report no evidence for the role of conduction delays in producing networks corresponding to MEG-derived networks of phase-synchronization. Further, we demonstrate the contribution of the topological organisation of the structural connectome and the dynamics of Wilson-Cowan oscillators in producing model-generated networks of phase-synchronization that correspond to those observed in MEG resting-state. Future studies will extend these models and subject them to local perturbations to provide insight on biophysical substrates underlying observed networks in rest and task conditions.

## Acknowledgements

The authors gratefully acknowledge the Academy of Finland (NW: 321542, SK: 292334, 319264, MP: 253130, 256472, 281414, 296304, 266745, SP: 266402, 266745, 303933, 325404), Department of Science & Technology (DST), India and Sigrid Juselius Foundation, for providing funding for this project. The authors are particularly grateful to Prof. Sitabhra Sinha and Prof. Mark Woolrich, for their invaluable comments and feedback, which contributed to this work. The authors are also grateful to Alexander Aushev, Anton Mallasto, Anirudh Jain and Diego Mesquita for comments on preliminary drafts of the manuscript.

## Supplementary figures

**Figure S1.**
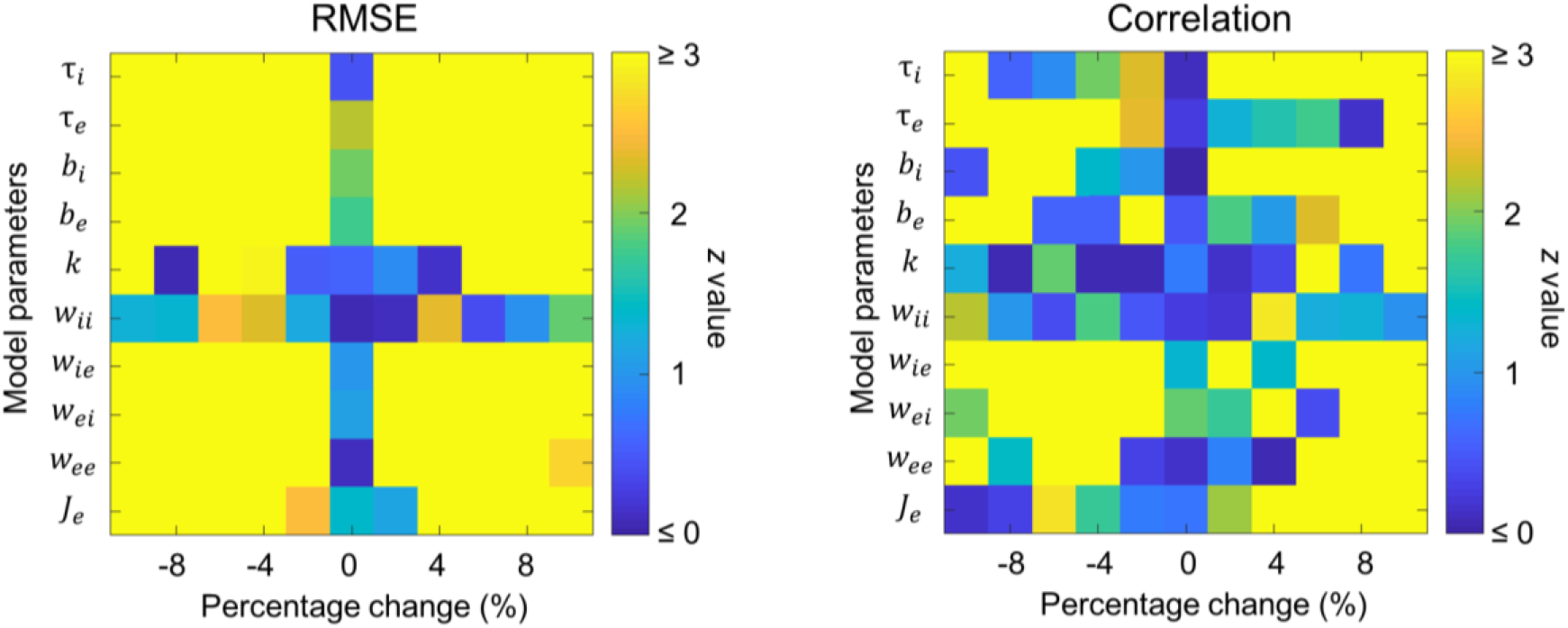
Model performance is sensitive to values of model parameters. Change in model performance, as measured by *z*-scored RMSE and *z*-scored Correlation, for percentage change in original value of each model parameter while keeping all other parameters at their original values.

**Figure S2.**
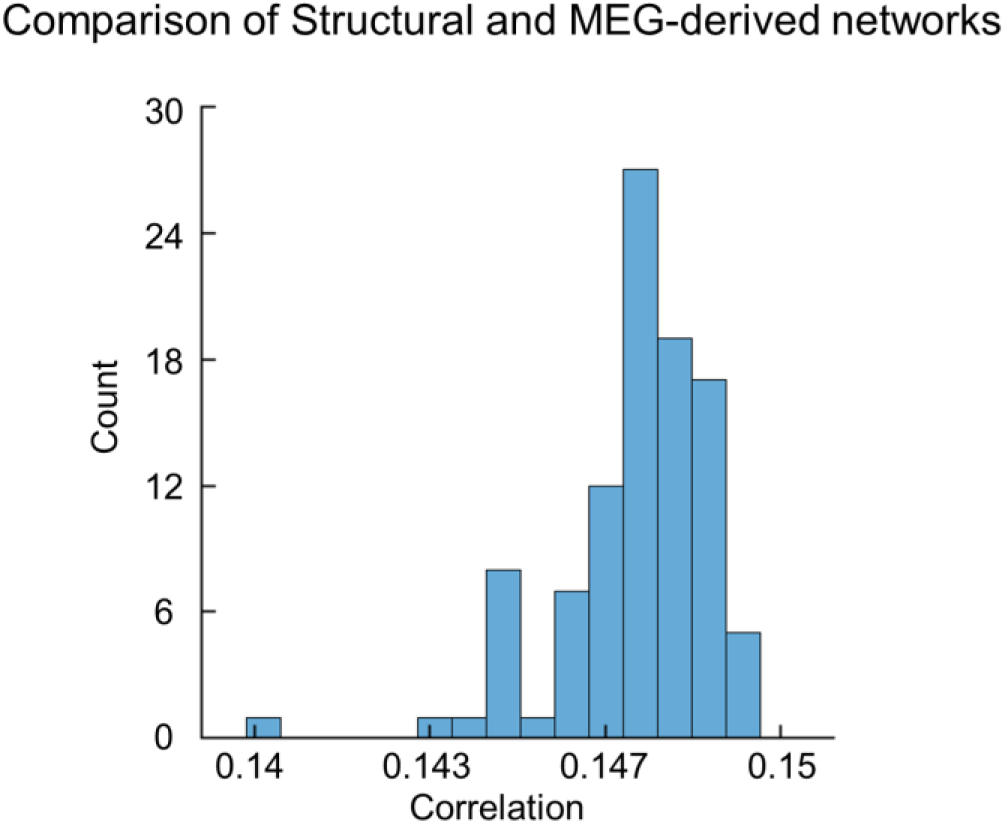
Structural connectome is weakly correlated to the MEG-derived network of phase-synchronization. Histogram of correlations between MEG-derived network of phase-synchronization and 100 bootstrapped versions of structural connectome.

**Figure S3.**
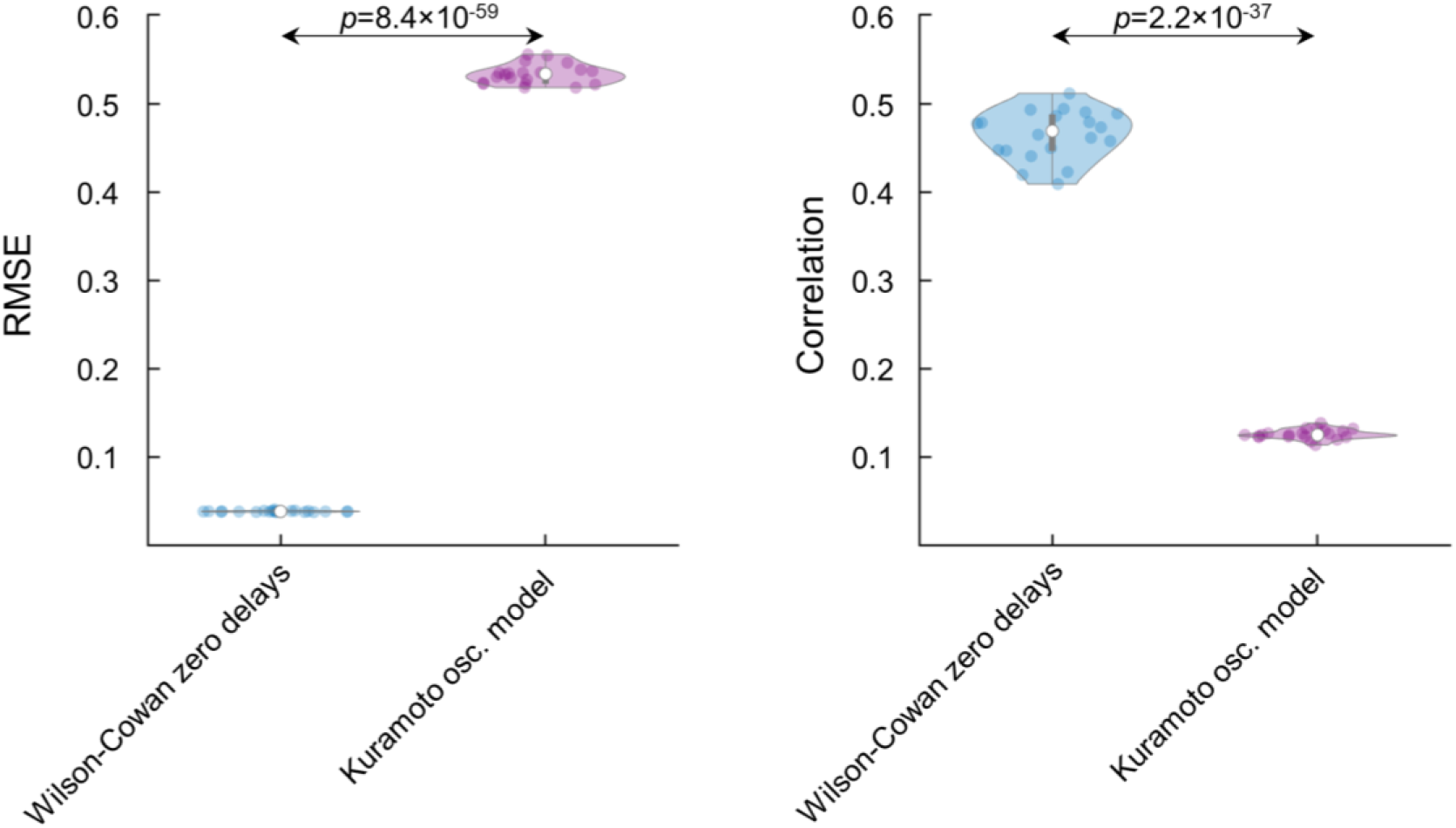
Wilson-Cowan zero delays model outperforms Kuramoto oscillator model. a. Violin plots of RMSE values for the Wilson-Cowan zero delays and Kuramoto oscillator models (left panel). Violin plots of Correlation values for the Wilson-Cowan zero delays and Kuramoto oscillator models (right panel).

**Figure S4.**
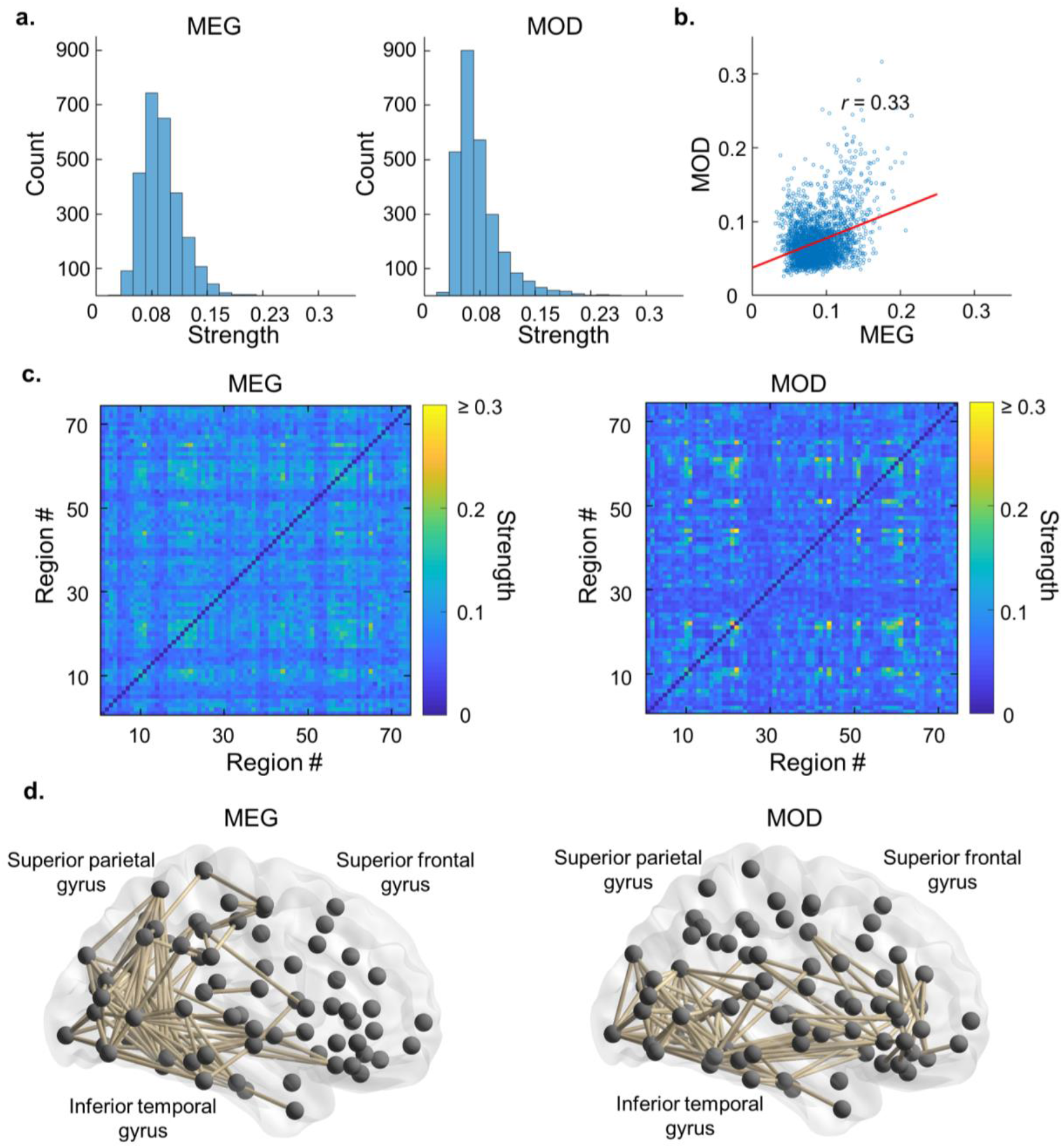
Moderate correspondence between mean model-generated network and right-hemispheric MEG-derived network of phase-synchronization. a. Histogram of connection strengths for mean model-generated and right-hemispheric MEG-derived networks of phase-synchronization b. Scatter plot of connection strengths for right-hemispheric MEG-derived network of phase-synchronization against mean model-generated network of phase-synchronization c. Matrices of connection strengths between every pair of brain regions for mean model-generated and right-hemispheric MEG-derived network of phase-synchronization. d. Brain network visualisation of top 5 percentile strongest connections for mean model-generated and right-hemispheric MEG-derived networks of phase-synchronization. MOD = model. Brain networks were visualised with BrainNet Viewer (http://www.nitrc.org/projects/bnv/) (Xia et al. (2013)).

